# Genome-Wide Transcriptome Dynamics in Auxin Homeostasis During Fruit Development in Strawberry (*F*. x *ananassa*)

**DOI:** 10.1101/2024.04.25.591171

**Authors:** Yoon Jeong Jang, Taehoon Kim, Makou Lin, Jeongim Kim, Kevin Begcy, Zhongchi Liu, Seonghee Lee

## Abstract

The plant hormone auxin plays a crucial role in regulating important functions in strawberry fruit development. Although a few studies have described the complex auxin biosynthetic and signaling pathway in wild diploid strawberry (*Fragaria vesca*), the molecular mechanisms underlying auxin biosynthesis and crosstalk in octoploid strawberry fruit development are not fully characterized. To address this knowledge gap, comprehensive transcriptomic analyses were conducted at different stages of fruit development and compared between the achene and receptacle to identify developmentally regulated auxin biosynthetic genes and transcription factors during the fruit ripening process. Similar to wild diploid strawberry, octoploid strawberry accumulates high levels of auxin in achene compared to receptacle. Consistently, genes functionating in auxin biosynthesis and conjugation, such as TRYPTOPHAN AMINOTRANSFERASE OF ARABIDOPSIS (TAAs), YUCCA (YUCs), and GRETCHEN HAGEN 3 (GH3s) were found to be primarily expressed in the achene, with low expression in the receptacle. Interestingly, several genes involved in auxin transport and signaling like PIN-FORMED (PINs), AUXIN/INDOLE-3-ACETIC ACID proteins (Aux/IAAs), TRANSPORT INHIBITOR RESPONSE 1 / AUXIN-SIGNALING F-BOX (TIR/AFBs) and AUXIN RESPONSE FACTOR (ARFs) were more abundantly expressed in the receptacle. Moreover, by examining DEGs and their transcriptional profiles across all six developmental stages, we identified key auxin-related genes co-clustered with transcription factors from the NAM-ATAF1,2-CUC2/ WRKYGQK motif (NAC/WYKY), BASIC REGION/ LEUCINE ZIPPER motif (bZIP), and APETALA2/Ethylene Responsive Factor (AP2/ERF) groups. These results elucidate the complex regulatory network of auxin biosynthesis and its intricate crosstalk within the achene and receptacle, enriching our understanding of fruit development in octoploid strawberries.

## Introduction

The garden strawberry (*Fragaria* ×*ananassa*), an allo-octoploid fruit crop (2n = 8x = 56), originated from the hybridization of two wild octoploid species, *F. chiloensis* subsp. *chiloensis* and *F. virginiana* subsp. *Virginiana*. It presents a genomic complexity with four subgenomes derived from different diploid progenitors: *F. vesca*, *F. iinumae*, *F. nipponica*, and *F. viridis*, as proposed by previous studies, (Edger et al., 2019; Hardigan et al., 2020). Strawberries belonging to the Rosaceae family, are distinguished by their unique achenetum fruit architecture, where a single flower produces many achenes (fertilized ovaries), contrasting with the fleshy fruits of apples, peaches, and pears (Xiang et al., 2017; Liu et al., 2020). This distinction in fruit type, notably in strawberries and raspberries as aggregate fruits significantly influences morphology, production, storage, and distribution of their unique fruits (Xiang et al., 2017; Liu et al., 2020; Veerappan et al., 2021). Fertilization-induced fruit development in strawberries is of interest due to their unique flower and fruit structure. The strawberry fruit consists of numerous individual achenes embedded in a fleshy receptacle (Tian et al., 2022). It is worth noting that what is commonly referred to as the fleshy fruit of a strawberry is actually a pseudocarp derived from the enlarged receptacle, while the true fruit (achene) is located on the epidermal layer (Perkins-Veazie, 1995; Guo et al., 2022). The fruit set is the primary and crucial point for fruit growth in plants, and typically triggered by positive signals generated during the process of fertilization (Gu et al., 1998). Fruit set can arise from different parts of the flower, such as the ovary in tomato (*Solanum lycopersicum*), receptacle in strawberry, and accessory part of hypanthium in apple (*Malus domestica*) (Gorguet et al., 2005; Pattison and Catalá, 2012; Kang et al., 2013; Galimba et al., 2019). In many plant species, the process of fruit set is largely dependent on the occurrence of fertilization. After successful pollination and fertilization, a series of physiological and molecular changes occur, ultimately leading to fruit development (Vivian-Smith et al., 2001). Flowers that do not undergo pollination and fertilization typically wither and fall off.

Plant growth hormones such as auxin, gibberellin, abscisic acid (ABA), and cytokinin, are essential regulators of strawberry fruit development. These hormones regulate diverse biological processes, including promoting growth, increasing fruit size, and enhancing fruit set and yield (Li et al., 2016; Liao et al., 2018; Katel et al., 2022). Particularly, auxin is essential for fruit development, as it regulates cell division and differentiation in the receptacle, and its levels are dramatically regulated during fruit growth and ripening (Feng et al., 2019). Although auxins such as indole-3-acetic acid (IAA) and phenylacetic acid (PAA) are crucial for plant growth and development, the complex biosynthetic pathways and enzyme redundancy within these pathways have hindered their complete understanding in plants, ranging from the model plant Arabidopsis to the crop model for fruit development (Woodward and Bartel, 2005; Mashiguchi et al., 2011). Previous studies have demonstrated that auxin transport in the achene ceased during the late stages of mid-green fruit development to the ripening process of strawberry. This leads to a decrease in auxin levels in the receptacle and the subsequent ripening process (Given et al., 1988). Furthermore, prior studies also reported that removal of the achene from the receptacle after pollination inhibits fruit enlargement, while exogenous application of auxin promotes receptacle growth in the absence of achene (Nitsch, 1950, 1952, 1955). Consequently, the achenes play a key role in auxin production needed to accelerate receptacle development (Nitsch, 1950). Despite numerous studies on the complex biosynthesis and signaling pathways of auxin in wild diploid strawberries (*F*. *vesca*), the fundamental molecular mechanisms of auxin biosynthesis in octoploid strawberries, based on the high-quality haplotype-phased reference genome, have not yet been elucidated. Additionally, the intricacies of the differential gene expressions between achenes and the receptacle during fruit development in octoploid strawberries still need to be characterized.

In Arabidopsis, IAA biosynthesis occurs mainly through the two-step biosynthetic pathways catalyzed by the amino transferases (Tryptophan aminotransferase of Arabidopsis (TAA) and TAA-related (TAR)), and monooxygenases belonging to the YUCCA (YUC) family that respectively convert Trp to Indole-3-pyruvic acid (IPyA) and IPyA to IAA (Mashiguchi et al., 2011; Won et al., 2011). Homologs of TAA/TAR and YUC have been identified in strawberry and implicated in various developmental processes, including fruit development. Several transcriptome profiling studies with *F. vesca* have shown that the auxin biosynthetic genes such as *FvYUC* and *FvTAA*, are predominantly induced in the endosperm after fertilization. Specifically, *FvYUC5*, *FvYUC11*, and *FvTAR1* were primarily expressed in the achene, with low expression in the receptacle, and highly expressed in ghost (seedcoat + endosperm) and less so in the embryo and ovary wall (Kang et al., 2013). It was also reported that *FvYUC4* and Fv*TAR2* were more abundantly expressed in the embryo, indicating their potential roles in embryonic development, while *FvYUC10*, *FvGA20ox1*, *FvGA20ox2*, and *FvGA3ox1* were present across various fruit tissues, exhibiting minimal to no expression within embryos. The expression of most of these biosynthetic genes gradually increases from stage 1 (open flower) to stages 5 (big green), likely due to the effect of fertilization (Kang et al., 2013). Additionally, it was discovered that the wild strawberry *F. vesca* possesses nine genetic loci of *YUCs* genes, eight of which encode functional proteins. Specifically, the overexpression of *FvYUC6* exhibited delayed flowering and male sterility in transgenic strawberry plants. Plants with reduced expression of *FvYUC6* through RNAi demonstrated alterations in floral organ structure and root development (Liu et al., 2014). In the transcriptome analysis of cultivated strawberries, two *YUC* genes, *FaYUC1* and *FaYUC2* (homologous to *AtYUC6* and *AtYUC4*, respectively), were identified. *FaYUC1* and *FaYUC2* are highly expressed in the large green fruit stage. Notably, the expression level of *FaYUC2* is much higher compared to *FaYUC1* (Liu et al., 2012). During late-stage fruit development, *FaYUC2*, *FaTAR2*, and *FaTAA1* were highly expressed in fruit receptacle (Estrada-Johnson et al., 2017; Feng et al., 2019).

Research on the plant hormone auxin, especially its perception and transcriptional regulation, is a critical aspect of plant phytohormone studies. Previous studies have identified auxin receptors, including the F-box protein Transport Inhibitor Response 1 (TIR1), which facilitates the degradation of AUXIN/INDOLE-3-ACETIC ACID (Aux/IAA) transcriptional repressors (Dharmasiri et al., 2005). At low auxin levels, Aux/IAA proteins act as repressors for expressing target genes like Auxin Response Factors (ARF). However, when auxin levels increase, Aux/IAA proteins are degraded by the 26S proteasome, leading to release ARFs to activate auxin responses (Calderón Villalobos et al., 2012). In *F*. *vesca*, studies on Aux/IAA and ARF genes suggest that the perception and transcriptional regulation of auxin involve 21 *FvAux/IAA*s and 19 *FvARF*s genes (Kang et al., 2013). Furthermore, the expression of 19 *FaAux/IAA* genes was detected in the receptacle of cultivated strawberries. Most of these genes exhibited a consistent decrease in expression as the fruit transitioned from green to red stages. However, *FaAux/IAA14b* and *FaAux/IAA11* showed maximum expression during the turning red stage, followed by a decrease in the red stage. For *FaARF* genes, while most showed low levels of expression from green to red stages, *FaARF6a* was notably more highly expressed during this transition compared to other *FaARF* genes (Estrada-Johnson et al., 2017). Despite the large amount of evidence, a detailed molecular mechanism controlling auxin perception and gene regulation in octoploid strawberries, particularly within specific tissues such as achenes and receptacles remains unclear.

In this study, a genome-wide transcriptome analysis was performed with a recent complete haplotype-phased octoploid strawberry reference genome of ‘Royal Royce’ (FaRR1) at various fruit developmental stages. Our results showed specific genes differentially expressed in receptacle and achenes during fruit development in octoploid strawberry. This deep transcriptome profiling identified genes involved in auxin biosynthetic signaling and metabolism pathways in strawberry. Moreover, our study identified novel transcription factors intricately linked to the auxin signaling network, marking a significant advancement in understanding the complex interplay between achenes and receptacles during the fruit development phase.

## Results

### Accumulation of IAA in achene and receptacle during fruit development

To investigate the developmental variations in IAA content of achenes and receptacles during fruit maturation, fruits of six different developmental stages were characterized. Stage 1; Small Green (SG), Stage 2; Medium Green (MG), Stage 3; Large Green (LG), Stage 4; White (W), Stage 5; Turning Red (TR), and Stage 6; Red (R) (Figure 1A). For each developmental stage, achenes and receptacles of individual fruits were separated and the concentration of IAA was quantified. The IAA content was consistently higher in achenes of all six stages, compared to the IAA detected in receptacles (Figure 1 B and C). The highest IAA content was observed in the achenes of the SG and MG stage (approximately 4,000 – 7,000 pmol/g FW). The concentration of IAA in achenes decreased as the fruit matured and showed the lowest level of IAA at stage R (approximately 1000 – 2000 pmol/g FW) (Figure 1B). Based on one-way ANOVA Turkey’s test, achenes at stages SG and MG contained significantly greater IAA content compared to the R stage (P < 0.05). In contrast, receptacles exhibited significantly reduced IAA content in all stages of fruit development, but the highest IAA level was seen in the SG stage (approximately 50-110 pmol/g FW). A rapid decrease in IAA content at the MG stage of receptacles was observed, with concentrations dropping below 20 pmol/g FW (Figure 1C). This reduction in IAA content in the receptacles after the SG stage persisted to maturity at the R stage.

**Figure 1.**
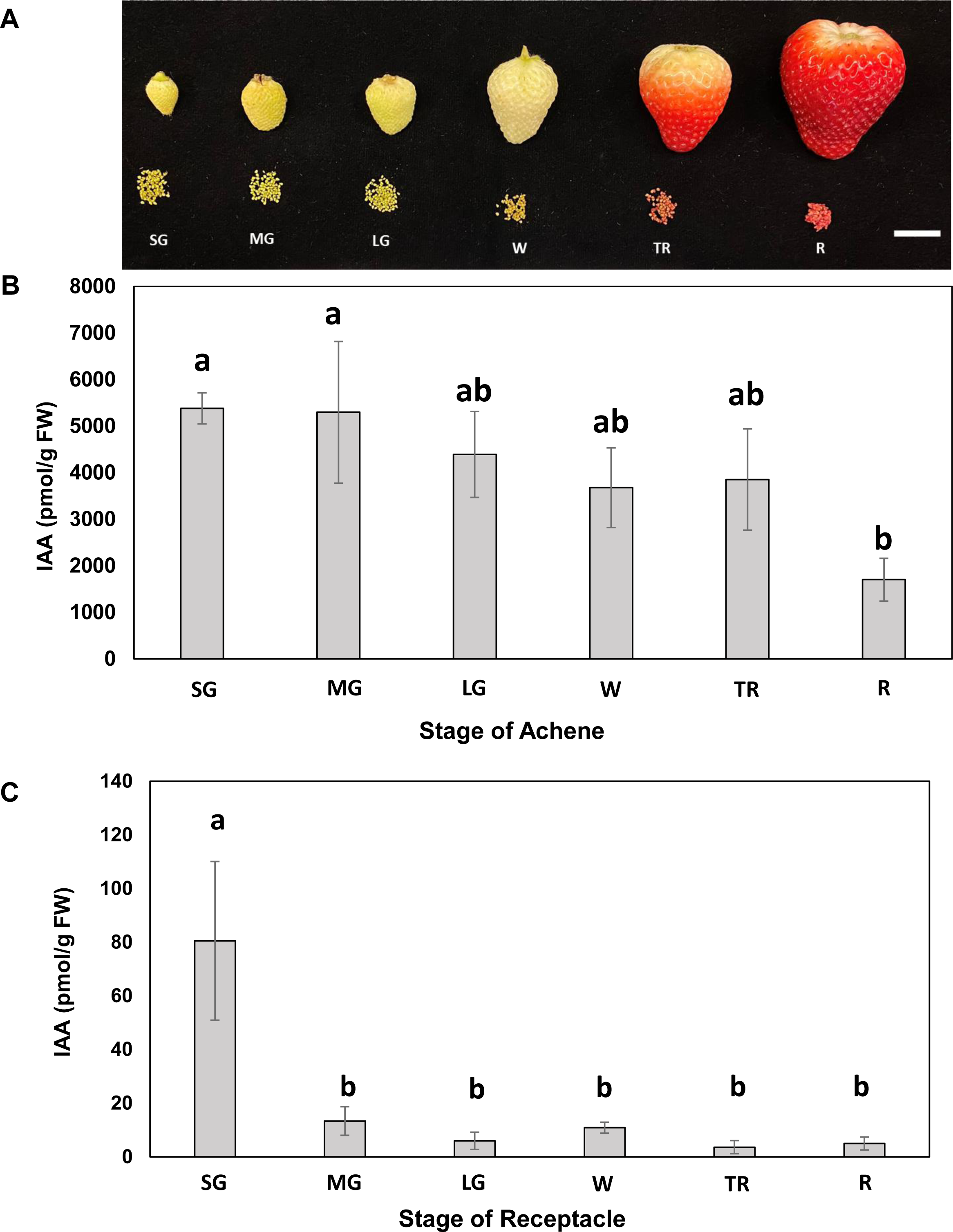
IAA levels in strawberry achene and receptacle during fruit development. A) Attached and detached achene and receptacle developmental stages of cultivar ‘Florida Brilliance’ (*Fragaria* × *ananassa* Duch). Stage 1; Small Green (SG), Stage 2; Medium Green (MG), Stage 3; Large Green (LG), Stage 4; White (W), Stage 5; Turning Red (TR), and Stage 6; Red (R). Scale bar: 2cm. IAA contents in (B) achenes and (C) receptacles. Statistical analysis was conducted using one-way ANOVA Tukey’s test (P < 0.05). Different letters indicate significant differences in the amount of free IAA. Mean and standard deviation are shown (n=3).

### Comparative transcriptome profiling analysis between achene and receptacle during fruit development

To investigate the transcriptional changes during achene and receptacle development (SG, MG, LG, W, TR, and R) related to auxin homeostasis an RNA sequencing approach was used. Sequencing reads were mapped to the octoploid strawberry reference genome, cv. ‘Royal Royce’ (FaRR1) (www.rosaceae.org), resulting in a high average mapping rate of 95.6% for achene samples and 95.3% for receptacle samples (Supplemental Table S1 and S2). Principal component analysis (PCA) was performed for the data quality assessment and exploratory analysis. Principal component 1 (PC1) accounted for 70% of the total variance, clearly delineating between achene and receptacle samples. Furthermore, principal component 2 (PC2) accounted for 22% of the total variance, distinguishing between the early and late stages of development. The PCA results suggest that the differences between achene and receptacle samples may reveal significant distinctions in the transcriptomes of these tissues across various stages of development (Supplemental Figure S1). To identify differentially expressed transcripts between achene and receptacle, we performed DESeq2 analysis using a significance cut-off padj <0.05. The larger number of DEGs was identified in the comparison between achene and receptacle at the MG stage (32,686 DEGs total; 17,608 up-regulated and 15,078 down-regulated). In contrast, the comparison between achene and receptacle at the R stage yielded the smallest number of DEGs, totaling 7,131 DEGs from which 6,217 were up-regulated and 914 were down-regulated (Figure 2A, Supplemental Figure S2 and Table S3). Venn diagrams were generated for pairwise comparisons between achene and receptacle at each stage (SG, MG, LG, W, TR, and R), and a total of 2,892 genes were found to be expressed in all six developmental stages (Figure 2A). Moreover, stage-specific analyses revealed that 3,612, 4,396, 3,069, 1,583, 817, and 189 genes were uniquely expressed at SG, MG, LG, W, TR and R stages, respectively (Figure 2A). Intriguingly, our findings reveal a notable decrease in the number of differentially expressed genes, particularly during the later developmental stages, including the W, TR, and R stages. Our results also showed a significant increase in the expression levels of achene-specific transcripts across the different fruit developmental stages. Additionally, the expression patterns of achene or receptacle-specific genes varied throughout different developmental stages (Figure 2B). These observations highlight the substantial expression of achene-specific transcripts during fruit maturation. The observed variation in gene expression patterns between achenes and receptacles during fruit maturation can be attributed to their distinct tissue compositions. Achenes, as more complex organs consisting of the embryo, endosperm, and seed coat, naturally display greater complexity and a higher number of expressed genes. Conversely, the receptacle, a simpler organ composed of cortex and pith, shows less variability in gene expression due to its more uniform tissue structure.

**Figure 2.**
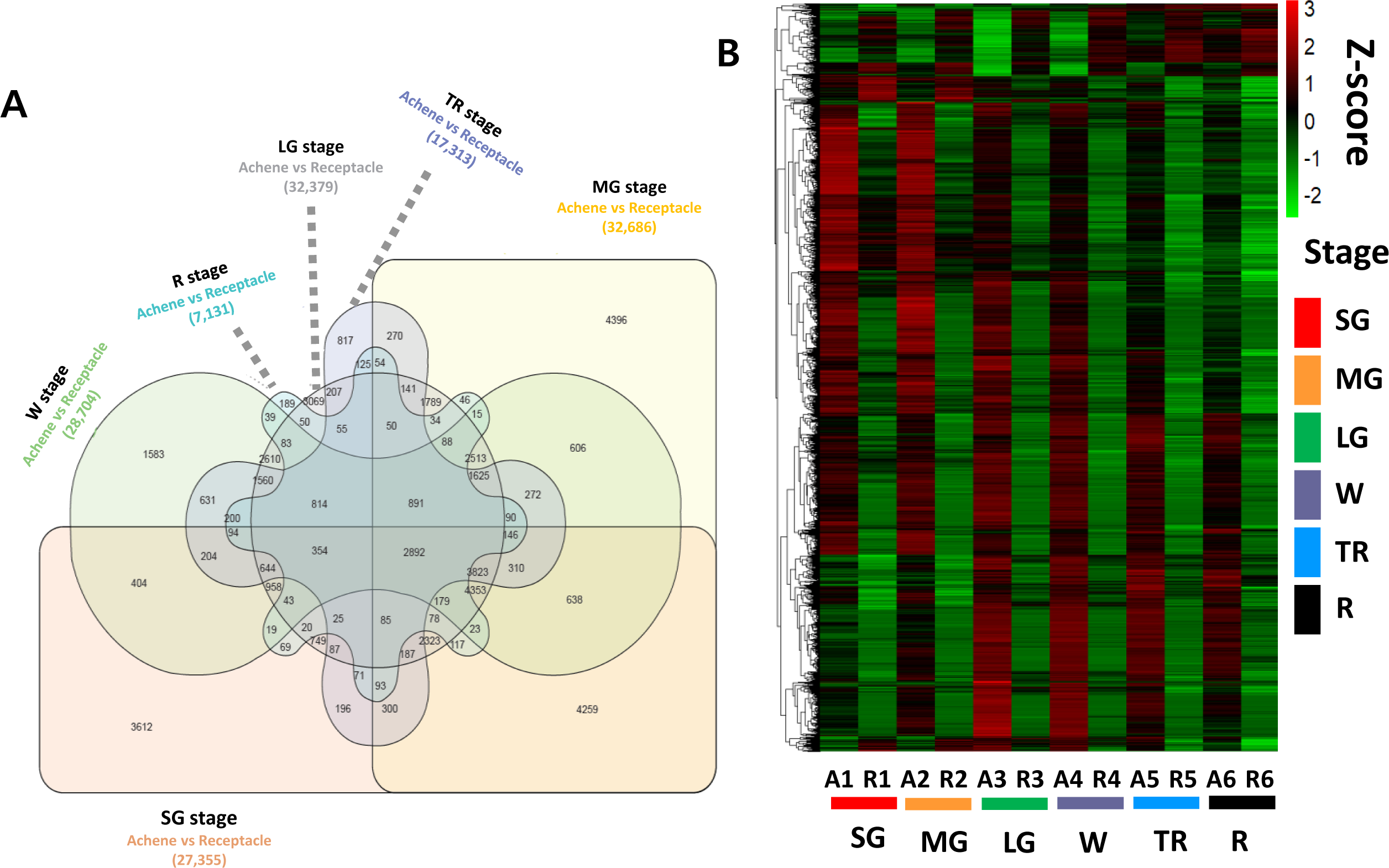
Transcriptional changes across six stages of fruit development in achene and receptacle tissues. A) Venn diagram displaying unique and overlapping sets of 2,892 DEGs identified in achene and receptacle tissues, padj < 0.05. B). Heatmap analysis of 2,892 overlapping DEGs. Gene expression values are shown as Z-score, padj < 0.05. Developmental stages are color coded. Small Green (SG) in red. Medium Green (MG) in yellow. Large Green (LG) in green. White (W) in purple. Turning Red (TR) in blue and Red (R) stage in black.

### Dynamic differential gene expression in achene and receptacle during fruit development

To elucidate the functional dynamics of gene expression between achene and receptacle tissues across six developmental stages, we conducted Gene Ontology (GO) and Kyoto Encyclopedia of Genes and Genomes (KEGG) pathway enrichment analyses using the ShinyGO web tool. We used a total of 2,892 DEGs identified across all different developmental stages of achene and receptacle (padj < 0.05) and subjected them to GO analysis to determine their associated Biological Process (BP), Cell Components (CC), and Molecular Functions (MF) associated with each gene (Figure 3 and Supplemental Table S4). Within the BP category, DEGs common to each of the six developmental stages in both achene and receptacle were primarily involved in pathways including the phenylpropanoid metabolic process, post-embryonic development, phenylpropanoid biosynthesis, as well as lignin metabolism and biosynthesis. Of particular note, we observed a common expression pattern of genes related to the response to hormone (GO:0009725) across all six stages (Figure 3, Supplemental Table S4). Additionally, in the CC category, the DEGs were found to relate to cellular components such as the vacuole, bounding membrane of organelle, vacuolar membrane, plant-type vacuole, and Golgi apparatus, among others (Figure 3, Supplemental Table S4). Interestingly, in the MF category, a significant enrichment of transcription factors and DNA binding activities was observed, including DNA-binding transcription factor activity, transcription regulator activity, transcription cis-regulatory region binding, transcription regulatory region nucleic acid binding, and sequence-specific DNA binding (Figure 3, Supplemental Table S4). Taken together, our findings suggest a conservation of transcriptional regulation mechanisms in the DEGs of achene and receptacle in octoploid strawberry. We also performed a KEGG pathway enrichment analysis on the 2,892 overlapped stage candidate DEGs identified. The primary signaling pathways associated with strawberry in the comparison between achene and receptacle include: Biosynthesis of secondary metabolites (139), Phenylpropanoid biosynthesis (31), Plant hormone signal transduction (32), Cutin, suberine and wax biosynthesis (9), stilbenoid, diarylheptanoid, and gingerol biosynthesis (4), Diterpenoid biosynthesis (6), Fatty acid degradation (9), and Flavonoid biosynthesis (6) (Figure 3B and Supplemental Table S5).

**Figure 3.**
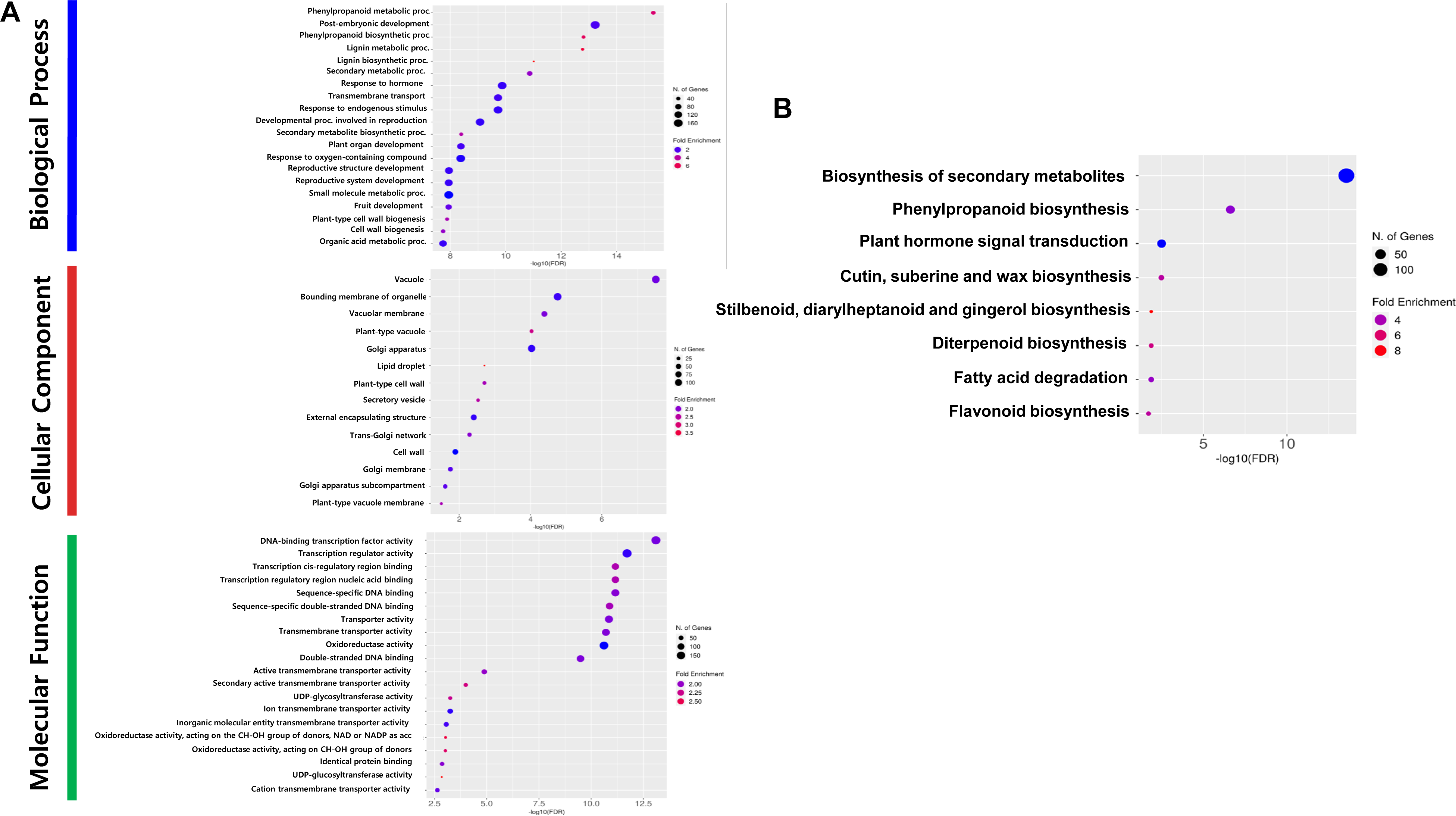
GO and KEGG pathway enrichment analyses of differentially expressed genes in achene and receptacle. A) Fold enrichment for GO biological process (BP), cellular component (CC) and Molecular function (MF). The x-axis indicates the -log10 FDR, and the y-axis shows the GO terms. Colors represent the degree of enrichment. B) Bubble diagram highlights eight significant KEGG pathways, with circle sizes reflecting the number of DEG counts. Pathways with the highest significance (FDR < 0.05) are delineated within respective clusters.

### Identification of genes involved in auxin homeostasis in achene and receptacle during fruit development of octoploid strawberry

The expression levels of genes associated with auxin homeostasis were determined in achene and receptacle, and a total of 164 genes were identified (Figure 4, Supplemental Table S6). The TAAs and YUCs are essential components for the biosynthesis of the major natural auxin in plants, IAA. As the IPyA pathway mediated by TAAs and YUCs produces the majority of free IAA, the expression of TAAs and YUCs is indispensable for various essential developmental processes, including fruit development. In the octoploid strawberry ‘Royal Royce’, we identified *TAA1* homoeologous gene copies located in the four subgenome of chromosome 4, *Fxa4Ag102498, Fxa4Bg102453, Fxa4Cg202161*, and *Fxa4Dg101990* (Figure 4). Among these copies, we found that *Fxa4Bg102453, Fxa4Cg202161*, and *Fxa4Dg101990* had high expression levels at the SG stage of achenes. Moreover, we identified homoeologous gene copies of *TAR1/2*, including *Fxa5Ag200524*, *Fxa5Bg100503*, *Fxa5Cg200481*, and *Fxa5Dg200494*, respectively (Figure 4). All four copies showed high expression levels only during the SG and MG stages. For *TAR3/4*, four homoeologous gene copies, *Fxa2Ag101303*, *Fxa2Bg201139*, *Fxa2Cg202166*, and *Fxa2Dg201128* were identified. Interestingly, only both *Fxa2Cg202166* and *Fxa2Dg201128* were highly expressed in the receptacle, particularly during the MG stage, compared to the achene (Figure 4).

**Figure 4.**
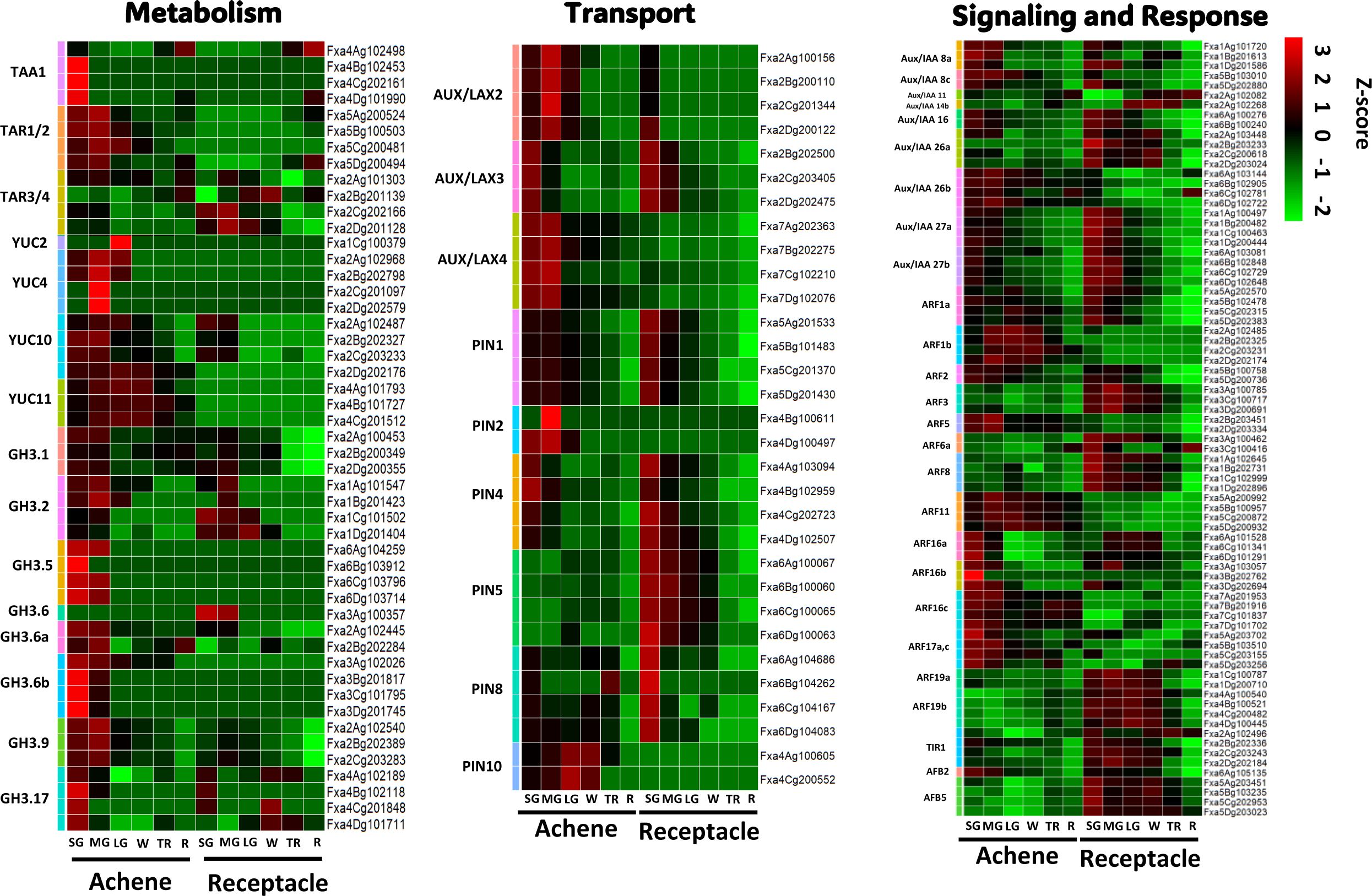
Transcriptional analysis reveals differential gene expression patterns of auxin metabolism, transport, signaling, and response genes in achene and receptacle during fruit development. Gene expression levels were normalized to z-scores using a log2(1+TPM) transformation and visualized using a color scale heatmap plot. Color gradients ranging from green to red represent the respective gene expression values, with green indicating lower expression and red indicating higher expression. Small Green (SG), Medium Green (MG), Large Green (LG), White (W), Turning Red (TR) and Red (R) stages.

In addition, four *FaYUC* genes, *FaYUC2*, *FaYUC4*, *FaYUC10* and *FaYUC11*, were found in the octoploid strawberry. Protein analysis using the *FvYUC2* sequence from diploid strawberry showed high similarity with four homoeologous copies of *FaYUC2*, *Fxa1Ag100425, Fxa1Bg200419, Fxa1Cg100379*, and *Fxa1Dg200387* (Figure 4). The gene expression result showed that only *Fxa1Cg100379* appears to be dominantly expressed at the LG stage at achene, while the expression of the other three copies is either absent or very low. For *FaYUC4*, interestingly, all four homoeologous copies of *Fxa2Ag102968, Fxa2Bg202798, Fxa2Cg201097*, and *Fxa2Dg202579* located in chromosome 2 are highly expressed during the MG stage of achene development. However, the expression of *FaYUC4* was only found at the subgenome A (*Fxa2Ag102968*), and B (*Fxa2Bg202798*) at the SG and LG stages. The gene *FaYUC10* has four homoeologous copies, *Fxa2Ag102487*, *Fxa2Bg202327*, *Fxa2Cg203233* and *Fxa2Dg202176*, in all subgenomes of chromosome 2. The expression levels were constitutively higher during the SG, MG, and LG stages of the achene compared to other stages. In receptacle, *Fxa2Ag102487*, *Fxa2Bg202327*, and *Fxa2Cg203233* show relatively higher expression levels at the SG and MG stages. For *FaYUC11*, three homoeologous gene copies, *Fxa4Ag101793*, *Fxa4Bg101727*, and *Fxa4Cg201512*, were found and showed higher expression in all developmental stages of the achene compared to the receptacle (Figure 4).

Furthermore, we identified high expressions of *GH3* genes, which belong to one of the major auxin-responsive gene families. *FaGH3.1*, *FaGH3.5*, *FaGH3.6a*, *FaGH3.6b*, *FaGH3.9*, and *FaGH3.17* showed expression levels with LogFC values from 0.48 to 6.06 (average= 2.4) at the SG stage and from 0.57 to 7.97 (average= 2.67) at the MG stage in achenes, respectively (Figure 4, Supplemental Table S3 and S6). It has been known that *GH3s* play crucial roles in auxin homeostasis through conjugating free auxin with amino acids (Wang et al., 2008). We identified that the *FaGH3.5* homoeologous gene copies, *Fxa6Ag104259*, *Fxa6Bg103912*, *Fxa6Cg103796*, and *Fxa6Dg103714*, were highly expressed in the achene during the SG and MG stages. Interestingly, only one copy of *FaGH3.6* (*Fxa3Ag100357*) was highly expressed in receptacle during the SG and MG stages compared to the achene. Furthermore, *FaGH3.6a*, *FaGH3.6b*, *FaGH3.9*, and *FaGH3.17* showed predominantly high expression levels at the SG and MG stages of achene. Particularly, the three homoeologous copies of *FaGH3.6b*, *Fxa3Bg201817*, *Fxa3Cg101795*, and *Fxa3Dg201745*, showed high expression levels in the achene at the SG stage (Figure 4).

For the PIN family genes, six homologs of PIN 1, 2, 4, 5, 8, and 10 were identified in *F. × ananassa* utilizing *F. vesca* as a query. Among them, *FaPIN1*, *FaPIN4*, *FaPIN5*, and *FaPIN8* exhibited stronger expression in the receptacle at SG than in achene at SG, but *FaPIN10* showed the opposite pattern. Homologs of AUX/LAX genes, including *AUX/LAX2*, *AUX/LAX3*, and *AUX/LAX4*, were identified in *F. × ananassa*. While *FaAUX/LAX2* exhibited high gene expression at MG of achene. *FaAUX/LAX4* showed stronger gene expression at SG and MG of achene compared to receptacle (Figure 4). We also discovered nine *Aux/IAA* genes and 14 ARF gene family homologs in *F. × ananassa*. Among them, *FaAux/IAA26b* displayed differences in gene expression between achene and receptacle at MG. *FaAux/IAA27a* and *FaAux/IAA27b* showed slight differences in expression between achene and receptacle at SG and MG. Notably, at SG and MG, *FaARF3*, *FaARF6a*, *FaARF19a*, and *FaARF19b* gene families exhibited higher expression in receptacle than in achene. However, *FaARF1b* displayed higher expression in achene from MG to W, and some genes including *FaARF16a*, *FaARF16b*, and *FaARF17a,c* showed higher expression in SG of achene compared to receptacle. Lastly, TIR1, AFB2, and AFB5 homologs were also identified in *F. × ananassa*. These auxin biosynthesis-related genes influence various aspects of *F. × ananassa* fruit effects, including growth, perception of auxin, and more, throughout the stages from early to late, suggesting their multifaceted impact on the fruit development of octoploid strawberry (Figure 4).

### Transcription factors involved in auxin biosynthesis signaling networks between achenes and receptacles during fruit development

To further understand auxin biosynthesis and transport between achene and receptacle, we used the entire list of DEG and clustered them across all six developmental stages based on their transcriptional expression. We identified four distinctive patterns of expression (Figure 5, Supplemental Table S7). Cluster 1 was formed by 1209 genes with increasing expression in achene having the highest expression at LG. Genes in cluster 1 have a low or no expression in the six receptacle developmental stages (Figure 5A). Cluster 2 formed by 1229 genes showed a decreased expression from achene to receptacle in a developmental manner. The more advanced the developmental stage the lower the expression (Figure 5A). Cluster 3 showed an opposite pattern as cluster 1, having lower expression in the achene and higher expression in the receptacle (Figure 5A). Finally, cluster 4 formed by 177 genes, showed variable expression across developmental stages in achene and receptacles samples (Figure 5A).

**Figure 5.**
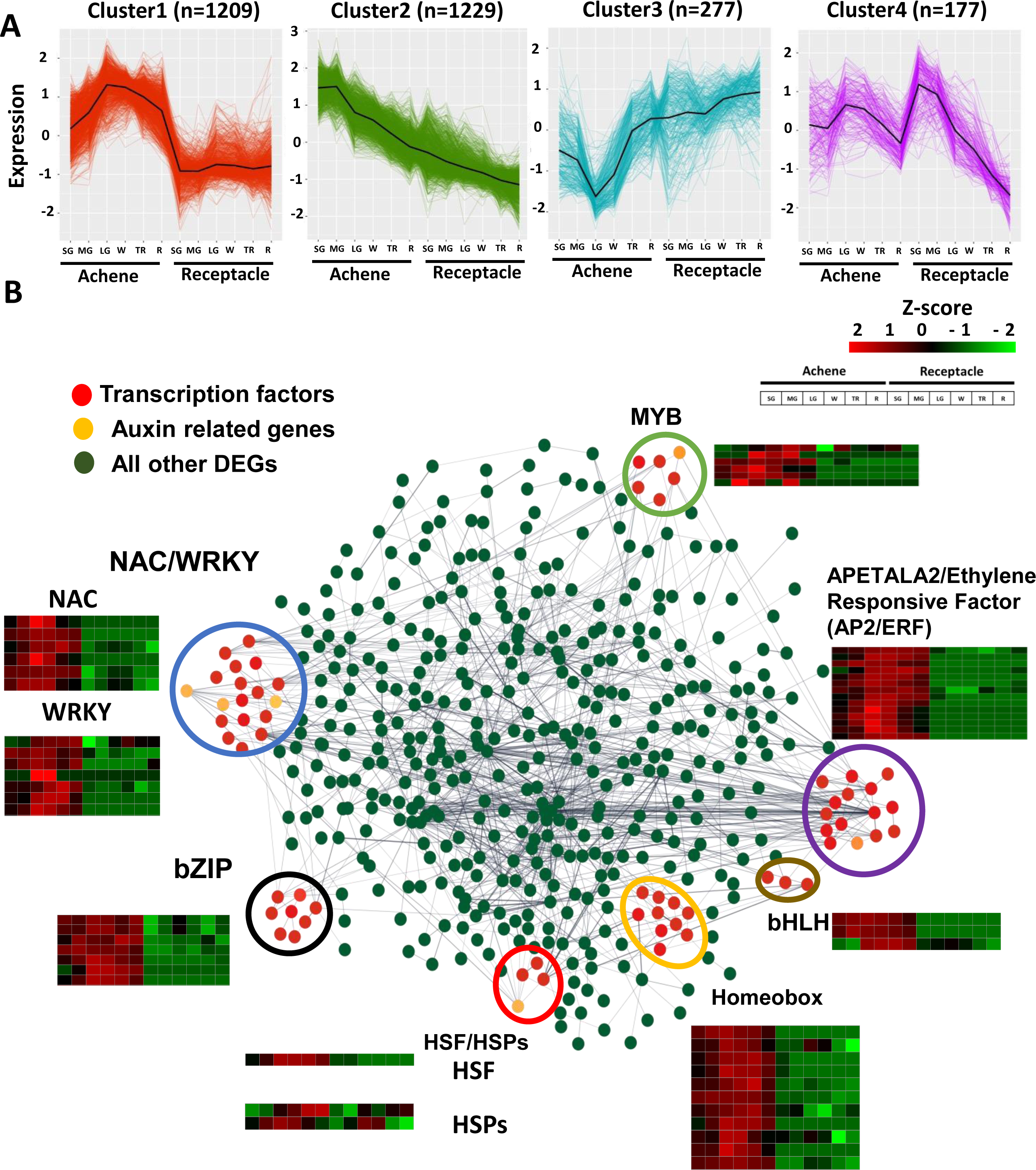
Novel transcription factors involved in gene regulation between achenes and receptacles of strawberry fruits. A) Clustering analysis including all differentially expressed genes. B) Gene network analysis of differentially expressed genes between achenes and receptacles. Transcription factors are shown in red. Auxin related genes are shown in yellow. All other differentially expressed genes are shown in green. A threshold of 0.7 of edge confidence was used. A detailed list of the genes included in the gene interaction analysis can be found in supplemental Table S7.

**Figure 6.**
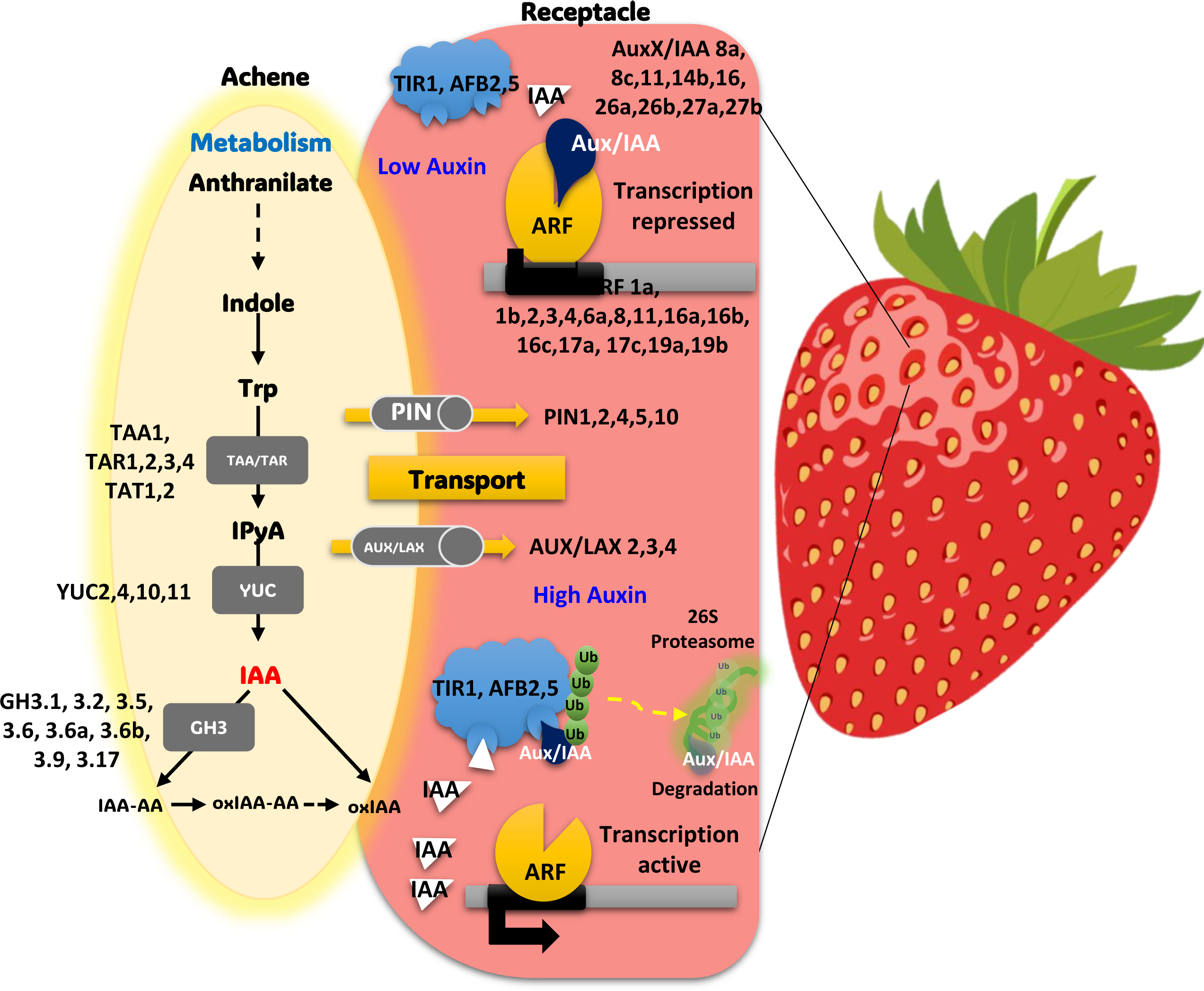
Model of auxin-related genes transcriptional behavior in achene and receptable of strawberry fruits. Auxin signaling pathway in strawberry development, highlighting key genetic components. Metabolic processes convert anthranilate to tryptophan, involving genes like TAA1 and TAT1,2, which lead to the production of the hormone auxin (IAA). The transport of auxin is facilitated by PIN and AUX/LAX proteins, crucial for establishing concentration gradients within the plant. In response to auxin levels, the Aux/IAA proteins may regulate gene expression by either repressing or permitting the activity of ARF proteins. High auxin levels trigger the degradation of Aux/IAA repressors, allowing ARF to activate transcription. Genes such as TIR1 and AFB2,5 are essential for auxin perception, initiating the proteasomal degradation pathway.

Since cluster 1 showed opposite developmental expression between achene and receptacle, we selected the entire set of genes identified in this cluster to conduct a correlation analysis to pinpoint transcription factors with shared expression profiles with genes involved in auxin biosynthesis and transport between achene and receptacle. A large portion of the strawberry genes are described as uncharacterized or unknown. Therefore, we removed them from the analysis. After removal of unrooted genes, the remaining genes were clustered using an edge confidence of at least 0.7. Our analysis yielded six main hubs containing transcription factors associated with auxin related genes (Figure 5B). These transcription factor hubs were grouped into NAC/WYKR (blue), MYB (green), APETALA2/Ethylene Responsive Factor (purple), homeobox (orange), heat shock transcription factor (red) and bZIP (black). Since we were interested in auxin biosynthesis and transport, we identified the auxin related genes (yellow) that clustered together with other transcription factors. Three out of the six hubs contained auxin related genes and were clustered with the NAC/WYKR, bZIP and APETALA2/Ethylene Responsive Factor hubs (Figure 5B). *FvYUC10* (*Fxa2Ag102487*) and *FaPIN10* (*Fxa4Ag100605*) clustered together with the APETALA2/Ethylene Responsive Factor group. *FvARF1a* (*Fxa5Dg202383*) and *FvARF16a* (*Fxa6Dg101291*) co-expressed with the NAC/WYKR hub and *FvARF11* (*Fxa5Dg200932*) with the bZIP group of transcription factor. Taken together, our analysis identifies potential transcription factors involved in the regulation of auxin and transport between achene and receptacles.

## Discussion

The plant hormone auxin plays a fundamental role in regulating cell expansion and cell division (Petersson et al., 2009; Vaddepalli et al., 2021). Previous studies indicate that fruit development is intricately linked to achene growth, with the maturation of fleshy fruits being inhibited in the absence of achene. This repression is relieved when fertilized achenes release auxin signals to the ovary wall and receptacle (Alabadí et al., 2009). In strawberries, achenes are considered the principal sites for auxin accumulation, playing a pivotal role in regulating auxin distribution across the entire fruit, thereby influencing its development. (Given et al., 1988; Kang et al., 2013; Sánchez-Sevilla et al., 2017). Studies on auxin in wild diploid strawberry, *Fragaria vesca*, have shown that auxin increases both the width and length of the fruit during the early stages of fruit development, while gibberellic acid (GA), mainly promotes vertical growth (Liao et al., 2018).

Perkins-Veazie classified the ripening stages of strawberry fruit into four stages (green, white, pink, and red), with pink being reported as the turning point of ripening (Perkins-Veazie, 1995). However, given that several previous studies have indicated the significant role of auxin synthesis and transport in the early developmental stages of strawberry growth, our research has not only divided the developmental process into six distinct stages, from early development to maturity but also separated achene and receptacle at each stage for hormone measurement and transcriptome analysis (Figure 1). A previous study has reported a consistent reduction in IAA as the fruit transitions from the small green stage to the red stage, as shown in mixed tissue of achene and receptacle (Symons et al., 2012). In this study, when analyzing the auxin content separately in the achene and receptacle of octoploid strawberries, we observed a significantly higher concentration of auxin in the achene compared to the receptacle, as observed in Fig 1. This suggests that receptacle enlargement is likely due to auxin moving from achenes.

Our results corroborate those of Estrada-Johnson et al. (2017) and Gu et al. (2019), who observed elevated IAA levels in achenes compared to receptacles during fruit development. Our study, however, distinguishes itself by employing a high-quality reference genome to perform detailed tissue-specific analyses across six developmental stages in octoploid strawberries, whereas Estrada-Johnson et al. (2017) focused on four stages. Moreover, while Gu et al. (2019) examined the transcriptome and auxin-related genes in diploid strawberries, our analysis extends to octoploid strawberry varieties, providing a more comprehensive understanding of auxin dynamics across different genomic complexities (Estrada-Johnson et al., 2017; Gu et al., 2019). One intriguing finding is the presence of approximately 50 - 110 pmol/g FW of IAA in the receptacles at the small green stage. This supports the result of temporal variations in auxin distribution, where auxin is synthesized within the achene and transported to the receptacle. Although present at a lower concentration compared to the achene, the IAA in the receptacle likely contributes to the coordination of fruit development, potentially regulating processes such as cell division and expansion. This supports the notion that even with a lower concentration, the receptacle serves as a receiver and transporter of IAA to modulate the growth and development of the strawberry fruit. To investigate the regulation of this phenomenon by specific genes, we conducted a genome-wide comprehensive analysis of the auxin-specific transcriptome in the achene and receptacle of octoploid strawberries using the recently developed high-quality haplotype-phased reference genome for octoploid strawberries (Hardigan et al., 2021). We compared and analyzed the transcriptomes separately for the achene and receptacle at each developmental stage. According to previous research, strawberry fruit set is completed within two to four days after fertilization, and it has been shown that transcriptional regulation and signaling metabolic changes occur at fertilization. This pattern is consistent with our results (Figure 1 and Figure 4) (Kang et al., 2013). To determine the collaborative action of gene families involved in the biosynthesis and transport of auxin, we conducted a DEG analysis using the criteria under padj < 0.05, log2(fold change) > 1 to identify significantly differentially expressed genes at each developmental stage.

In this study, we examined the expressions of homologous genes previously studied, such as TAA, TAR, YUC, GH3, Aux/IAA, TIR/AFB, and ARF, and found that the genes related to auxin biosynthesis were highly expressed during SG or MG in the achene. Throughout the comprehensive analysis in this study, it was found that genes such as TAA1, YUC4, and GH3.6b play a pivotal role in auxin biosynthesis primarily within the achene. Additionally, the PIN gene family, essential for auxin efflux facilitation, markedly impacts auxin’s directional flow owing to their unique intracellular localization (Petrásek et al., 2006; Adamowski and Friml, 2015). In octoploid strawberries, particularly during the SG stage, the expression of specific PIN genes (*FaPIN4*, *FaPIN5*, and *FaPIN8*) is substantially elevated in the receptacle compared to the achene. This reaffirms the coordination of achene and receptacle in fruit TAA development. Previous studies have also reported the high expression of auxin-conjugating GH3 genes in ghost and ovary walls at the *F. vesca* (Kang et al., 2013). Furthermore, the presence of IAA amide conjugates and highly abundant IAA-protein conjugates have been reported in the receptacle of strawberries (Archbold and Dennis, 1984; Park et al., 2006)

One notable aspect to consider is that in many species, a multitude of Aux/IAA proteins modulate ARF-mediated transcription and offer extensive signaling interactions in various processes involving auxin (Liscum and Reed, 2002). In this research, we discovered that gene copies within the Aux/IAA family, specifically *FaAux/IAA26a* (*Fxa2Ag103448*, *Fxa2Bg203233*, *Fxa2Cg200618*, *Fxa2Dg203024*), *FaAux/IAA27a* (Fxa1Ag100497, *Fxa1Bg200482*, Fxa1Cg100463, *Fxa1Dg200444*), and *FaAux/IAA27b* (*Fxa6Ag103081*, *Fxa6Bg102848*, *Fxa6Cg102729*, *Fxa6Dg102648*), alongside TIR/AFB genes such as *FaTIR1* (*Fxa2Ag102496*, *Fxa2Bg202336*, *Fxa2Cg203243*, *Fxa2Dg202184*), *FaAFB2* (*Fxa6Ag105135*), and *FaAFB5* (*Fxa5Ag203451*, *Fxa5Bg103235*, *Fxa5Cg202953*, *Fxa5Dg203023*), and ARF genes including *FaARF3* (*Fxa3Dg20069*, *Fxa3Cg100717*, *Fxa3Ag100785*), *FaARF6a* (*Fxa3Cg100416*, *Fxa3Ag100462*), *FaARF19a* (*Fxa1Cg100787*, *Fxa1Dg200710*), and *FaARF19b* (*Fxa4Ag100540*, *Fxa4Bg100521*, *Fxa4Cg200482*, *Fxa4Dg100445*), demonstrated notably higher expression levels in the receptacle than in the achene. This differential expression was particularly evident during the early developmental stages, from SG to LG, respectively. This intricate network of interactions may stem from gene expression driven by specific TIR/AFB, Aux/IAA, and ARF transcription factors in response to auxin (Weijers and Wagner, 2016). Furthermore, our study aimed to elucidate the transcription factors associated with the genes discovered during fruit development in octoploid strawberry. We focused on transcription factors that exhibit co-expression patterns with genes involved in auxin biosynthesis and transport within both achene and receptacle tissues. Our analysis revealed that several auxin-related genes were grouped alongside the NAC/WYKR, bZIP, and AP2/ERF transcription factor hubs. Specifically, *FaYUC10* (*Fxa2Ag102487*) and *FaPIN10* (*Fxa4Ag100605*) were associated with the AP2/ERF cluster. Auxin has been proposed to indirectly promote fruit ripening by stimulating the transcription of several ethylene components, leading to ethylene-induced fruit ripening and softening. Additionally, *FaARF1a* (*Fxa5Dg202383*) and *FaARF16a* (*Fxa6Dg101291*) demonstrated co-expression with the NAC/WYKR hub, while *FaARF11* (*Fxa5Dg200932*) aligned with the bZIP transcription factor group. This collective data suggests potential transcription factors that may play a crucial role in regulating auxin biosynthesis and its transport dynamics between achene and receptacle during strawberry fruit ripening.

In conclusion, our study provides comprehensive gene expression profiling specific at subgenome level of octoploid strawberries from early to late fruit developmental stages. This information highlights an important yet insufficiently explored area in the field of octoploid strawberry auxin research. Furthermore, identifying pattens of subgenome-specific gene expression would implicate pathways of auxin metabolites, as well as the transport and perception of auxin between achenes and receptacles. Leveraging recently available high-quality haplotype-phased reference genomes and genome-wide transcriptome profiling analysis, we were able to unveil the network of genes in auxin homeostasis and enhance our understanding of the regulatory mechanisms during fruit development in strawberry.

## Materials and methods

### Sample preparations

The samples of the strawberry fruits (*Fragaria* × *ananassa* Duch. cv. Brilliance) were used for transcriptome analysis. The six different developmental stages of harvested fruits corresponded that stage 1 is Small Green (SG), stage 2 is Medium Green (MG), stage 3 is Large Green (LG), stage 4 is White (W), stage 5 is Turning Red (TR), and stage 6 is Red (R). In early January, the fruits were harvested from the UF strawberry field at Gulf Coast Research and Education Center in Balm, Florida. All stages of the fruits, including transcriptome sequencing biological replicated, were harvested simultaneously. Using the forceps and scalpels, each stage of the achenes and receptacle were separated from the fruits.

### Auxin measurement

Approximately 20 mg (fresh weight) of achene and receptacle samples from 6 fruit stages were immediately frozen in liquid nitrogen and stored at -80°C. For IAA extraction and purification, frozen samples of achenes and receptacles were ground with pestles in liquid nitrogen and promptly submerged in 1.1 mL sodium phosphate buffer (50mM, pH 7.0) containing 0.1% diethyl dithiocarbamic acid sodium salt and 10 ng/mL of [^13^C_6_]-IAA as internal standard. Samples were incubated at 4°C with continuous shaking for 40 min and then centrifuged at 13000 *g* at 4°C for 15 min. After collecting supernatant and adjusting pH to 2.7, samples were purified by solid-phase extraction using Oasis^TM^ HLB columns (WAT094225; Waters, MA, USA). The final elutes with 80% methanol were evaporated to dryness *in vacuo* and stored at -20°C until LC/MS analysis.

The IAA detection method using liquid chromatography and mass spectrometer (LC-MS) was adapted from (Perez et al., 2021). All samples were resuspended in MilliQ water and analyzed using Vanquish Horizon ultra-high performance liquid chromatography (UHPLC) installed with an Eclipse Plus C18 column (2.1 × 50 mm, 1.8 μm) (Agilent) and mass analysis was performed using a TSQ Altis Triple Quadrupole (Thermo Scientific) MS/MS system with an ion funnel. MRM parameters of the standards (precursor m/z, fragment m/z, radio frequency (RF) lens and collision energy) of each compound were optimized on the machine using direct infusion of the authentic standards. IAA and [^13^C_6_]-IAA were purchased from Cambridge Isotope Laboratories. For IAA detection, the mass spectrometer was operated in positive ionization mode at an ion spray voltage of 4800 V. Formic acid (0.1%) in water and 100% acetonitrile were used as mobile phases A and B, respectively, with a gradient program (0–95% solvent B over 4 min) at a flow rate of 0.4 mL per min. The sheath gas, aux gas and sweep gas were set at 50, 9 and 1 (arbitrary units), respectively. Ion transfer tube temperature and vaporizer temperature were set at 325°C and 350°C, respectively. For MRM monitoring, both Q1 and Q3 resolutions were set at 0.7 FWHM with collision-induced dissociation (CID) gas at 1.5 mTorr. The scan cycle time was 0.8 s. MRM for IAA was used to monitor parent ion → product ion reactions for each analyte as follows: m/z 175.983→130.071 (CE, 18 V) for IAA; m/z 182.091→136 (CE, 18 V) for [^13^C_6_]-IAA. IAA analysis was conducted with three biological replicates.

### Transcriptome data analysis

The total RNA from separate achene and receptacle at six development stage with three replications per sample were extracted by the Spectrum^TM^ Plant Total RNA Kit (Sigma-Aldrich, MO, USA) as fallowing the manufacturer protocol. To generate RNA-seq for illumina sequencing library were fallowed Illumina sequencing protocol. The resulting sequencing library were performed pair-end sequenced (2 x 150bp) by Illumina NovaSeq instruments at Novogene Bioinformatics Institute, Beijing, China. Raw read sequences obtained from 36 sequenced libraries were quality trimmed and filtered using Trimmomatic (Bolger et al., 2014). Data quality was assessed using FastQC. Trimmed paired end reads were aligned to the ‘Royal Royce’ octoploid genome (Hardigan et al., 2021) using Hisat2 (Kim et al., 2019). Differentially expressed genes (DEG) analysis was conducted using the DESeq2 package in R script (Love et al., 2014) using a fold change >|1| and P < 0.05 (after the false discovery rate adjustment for multiple testing (FDR) <0.05) for the null hypothesis. Total number of reads mapped to each gene was used to calculate transcripts per million (TPM) values, which were determined using a custom Python script.

Gene Ontology (GO) enrichment analysis was conducted using Arabidopsis gene information provided by the Royal Royce genome annotation database (Hardigan et al., 2021). The analysis was conducted through the ShinyGO tool (version 0.77, http://bioinformatics.sdstate.edu/go/, (Ge et al., 2020)), applying a P-value cutoff of ≤ 0.05 (FDR) and default options.

### Confirmation of homology of octoploid strawberry auxin hormone pathway genes

Genes associated with auxin biosynthesis were explored in the octoploid strawberry involved utilizing known genes from *F. vesca* and previous literature. BlastP was performed with significant criteria under e-value=0, pident<90, bit score >100 based on the protein sequence against the Royal Royce reference genome (Li et al., 2019; Hardigan et al., 2021) for each hormone-related gene (Coulouris et al., 2008).

### K-means Clustering and Gene Co-expression Analysis

Transcripts per million (TPM) counts were used as input for the K-means clustering analysis. TPM values were averaged for the replicates of each tissue (achene or receptacle) at any given stage (Stage 1 to Stage 6). Averaged TPM values were normalized to log_2_(TPM+1), scaled, and used as input for clustering analysis. DEGs were categorized into four clusters using the k-means algorithm implemented using the R programming language. In order to identify transcriptional correlations among genes with shared expression profile analysis achene or receptacle, the *cor* function was implemented in the R package WGCNA. Then, genes with shared expression profiles were considered as seed candidates and were used to build a gene co-expression analysis to obtain direct and indirect interactions. A high confidence score of 0.7 was used as a threshold (Begcy et al., 2019). Only high levels of confidence interactions were considered as valid as used in the final gene co-expression analysis.

## Data and materials availability

The data underlying this article are available in Supplementary Information.

## Accession Number

High-throughput sequencing data analyzed in present study are available under NCBI BioProject PRJNA1010111.

## Supplementary Information

The online version contains supplementary material available

## Acknowledgements

We extend our gratitude to Dr. Vance M. Whitaker for providing the fruit materials used in this study. The authors thank for the technical support and assistance with fruit preparations provided by Dr. Youngjae Oh and Sadikshya Sharma. Also, we thank Ru Dai and Veronica Perez for their technical support for auxin quantification.

## Author contributions

YJ, KB, JK and SL contributed to the study design and drafted the article. YJ, JK, TK, KB, HH, ML, ZL, VW, SL and analyzed the experiment results, prepared figures and tables. All authors read and approved the manuscript.

## Declarations

### Supplementary Figs

**Figure S 1 Principal component analysis (PCA) of transcriptome data from 36 samples.** Each colored prototype demonstrates the distinct separation of developmental samples into different stages throughout the entire time course. Furthermore, the analysis includes 36 samples, encompassing different tissue types, namely achene and receptacle, as well as 3 biological replicates for each stage.

**Figure S 2 Comparison of vesicular and receptor differentially expressed genes via DESeq2.**

**Supplementary Tables**

**Table S1 Representative statistics RNA-Seq quality for transcriptome analysis of achene and receptacle samples.**

**Table S2 Individual RNA-Seq quality results for transcriptome statistics of achene and receptacle samples at each stage of fruit development.**

**Table S3 Transcript levels of differentially expressed genes in the six different stages of fruit development.**

**Table S4 List of GO pathway. Table S5 List of KEGG pathway.**

**Table S6 Transcriptomic levels of the differential gene expression patterns of auxin metabolism, transport, signaling, and response genes in achene and receptacle during developmental stages.**

**Table S7 List of novel transcription factors involved in gene regulation between achenes and receptacles of strawberry fruits.**

